# Identifying Dihydropyrimidine Dehydrogenase as a Novel Regulator of Hepatic Steatosis

**DOI:** 10.1101/2021.03.04.433987

**Authors:** Kelly E. Sullivan, Sheetal Kumar, Ye Zhang, Emily de Koning, Jing Yuan, Fan Fan

**Affiliations:** Amgen Inc. Cambridge, MA 02141

## Abstract

Pyrimidine catabolism is implicated in hepatic steatosis. Dihydropyrimidine Dehydrogenase (DPYD) is an enzyme responsible for uracil and thymine catabolism, and DPYD human genetic variability affects clinically observed toxicity following 5-Fluorouracil (5-FU) administration. In an *in vitro* model of diet-induced steatosis, the pharmacologic inhibition of DPYD resulted in protection from lipid accumulation. Additionally, a gain-of-function mutation of DPYD, created through clustered regularly interspaced short palindromic repeats associated protein 9 (CRISPR-Cas9) engineering, led to an increased lipid burden, which was associated with altered mitochondrial functionality in a hepatocarcionma cell line. The studies presented herein describe a novel role for DPYD in hepatocyte metabolic regulation as a modulator of hepatic steatosis.

## Introduction

Hepatic steatosis results from dysregulated energy homeostasis in the liver and an accumulation of lipids in the parenchyma (1). Hepatic steatosis, or fatty liver, is the first stage of the nonalcoholic fatty liver disease (NAFLD) spectrum. Some individuals with steatosis may progress to non-alcoholic steatohepatitis (NASH), the stage at which excessive levels of triglycerides become cytotoxic and trigger an inflammatory and subsequent fibrotic response (2). This disease process impairs normal hepatic function and is quickly becoming a global epidemic with nearly 30% of the population impacted (3). NASH is a precursor stage to cirrhosis, and cirrhosis from fatty liver is increasingly becoming the leading reason for liver transplantation globally (4). Currently there are no approved therapies for NASH or cirrhosis. Identifying novel protein targets of hepatic lipid accumulation is key to understanding and preventing disease pathogenesis (5).

Pyrimidine catabolism has been implicated in liver lipid homeostasis. Transgenic mouse models identify the enzyme, uridine phosphorylase-1 (UPP1), as a key driver of pyrimidine catabolism and the salvage pathway, and a modulator of hepatic microvesicular steatosis (6). An intermediate of pyrimidine catabolism, dihydrothymine, has been identified as a metabolic signature associated with pediatric NAFLD (7). Daily uridine supplementation for two weeks improved liver lipid metabolism and weight loss in mice fed a high fat diet (8) and reduced drug-induced liver lipid accumulation in mice following exposure to hepatotoxic drugs such as tamoxifen (9), zalcitabine (10) and fenofibrate (11). However, chronic uridine administration for 16 weeks induced fatty liver and glucose intolerance in mice (12). In addition, plasma uridine levels are regulated by fasting and feeding cycles in mice in an adipocyte dependent manner and uridine levels can directly influence energy homeostasis and thermoregulation (13). These results suggest pyrimidine catabolism regulates metabolism and sensitive pharmacologic control of uridine levels may be a promising approach to treating metabolic disorders.

Uridine catabolism is accomplished through its reversible conversion to uracil via the UPP1 enzyme (6). The enzyme dihydropyrimidine dehydrogenase (DPYD) is the rate limiting enzyme in uracil catabolism and therefore a major regulator of uracil and uridine levels in the liver (14). DPYD expression levels are controlled in part by the farsenoid X receptor (FXR) (15), a transcription factor known to regulate hepatic energy homeostasis. A high degree of genetic variability in DPYD has been highlighted in the human population through the development of the chemotherapeutic 5-Fluorouracil (5-FU) for the treatment of colorectal cancer. The majority of 5-FU is metabolized and cleared through the liver via DPYD, thus controlling the fraction of active metabolites present in the bloodstream. Loss-of-function carriers in DPYD demonstrate significant toxicity to 5-FU due to higher levels of circulating drug, whereby gain-of-function carriers demonstrate reduced responsiveness to treatment because of higher rates of clearance (16). In addition, pharmacological manipulation of DPYD through the small molecule inhibitor, Gimeracil, has, clinically, demonstrated the ability to increase the efficacy of 5-FU treatment (17).

Whether genetic variability and/or pharmacologic perturbation of DPYD alters uracil and uridine homeostasis, resulting in liver lipid accumulation and susceptibility to NAFLD, remains to be elucidated. Thus, these questions were explored using a variety of *in vitro* cell models. The results demonstrate that inhibition of DPYD results in a reduced lipid burden in the presence of free fatty acid exposure in primary human hepatocytes cultured *in vitro*. RNAseq and extracellular flux analysis revealed a reduction in lipogenesis and an increase in mitochondrial respiration likely due to increased beta-oxidation of fatty acids with exposure to the DPYD inhibitor, Gimeracil. A gain-of-function mutation in DPYD was engineered into a human hepatocarcinoma cell line using CRISPR-Cas9 homology-directed repair to assess whether increased DPYD activity altered the basal lipid profile of the cells and their response to stress. Mechanistic evaluation revealed a reduced lipid burden in a DPYD-knockout cell line and an increased lipid burden with the L310S homozygous gain of function mutation. Extracellular flux analysis demonstrated impaired mitochondrial respiration in the L310S line and improved functionality in the loss-of-function line. These human *in vitro* models of hepatic steatosis highlight how DPYD enzymatic activity significantly contributes to lipid homeostasis and demonstrate how pharmacologic inhibition of DPYD may be a potential treatment option for the mitigation of diet-induced microvesicular steatosis.

## Results

Following exposure to free fatty acids (FFAs), PHHs demonstrated a dose-dependent increase in neutral lipid accumulation (Figure 1A and 1B) and a significant increase in basal oxygen consumption rate as measured by extracellular flux analysis (Figure 1E). Significant disruptions in expression of mRNA transcripts associated with maintenance of metabolic homeostasis, including upregulation of *SLC25A47, APOA4, APOA5, MFSD2A*, and *ABCB4*, and downregulation of *ABCG1* were observed (Figure 1D).

**Figure 1.**
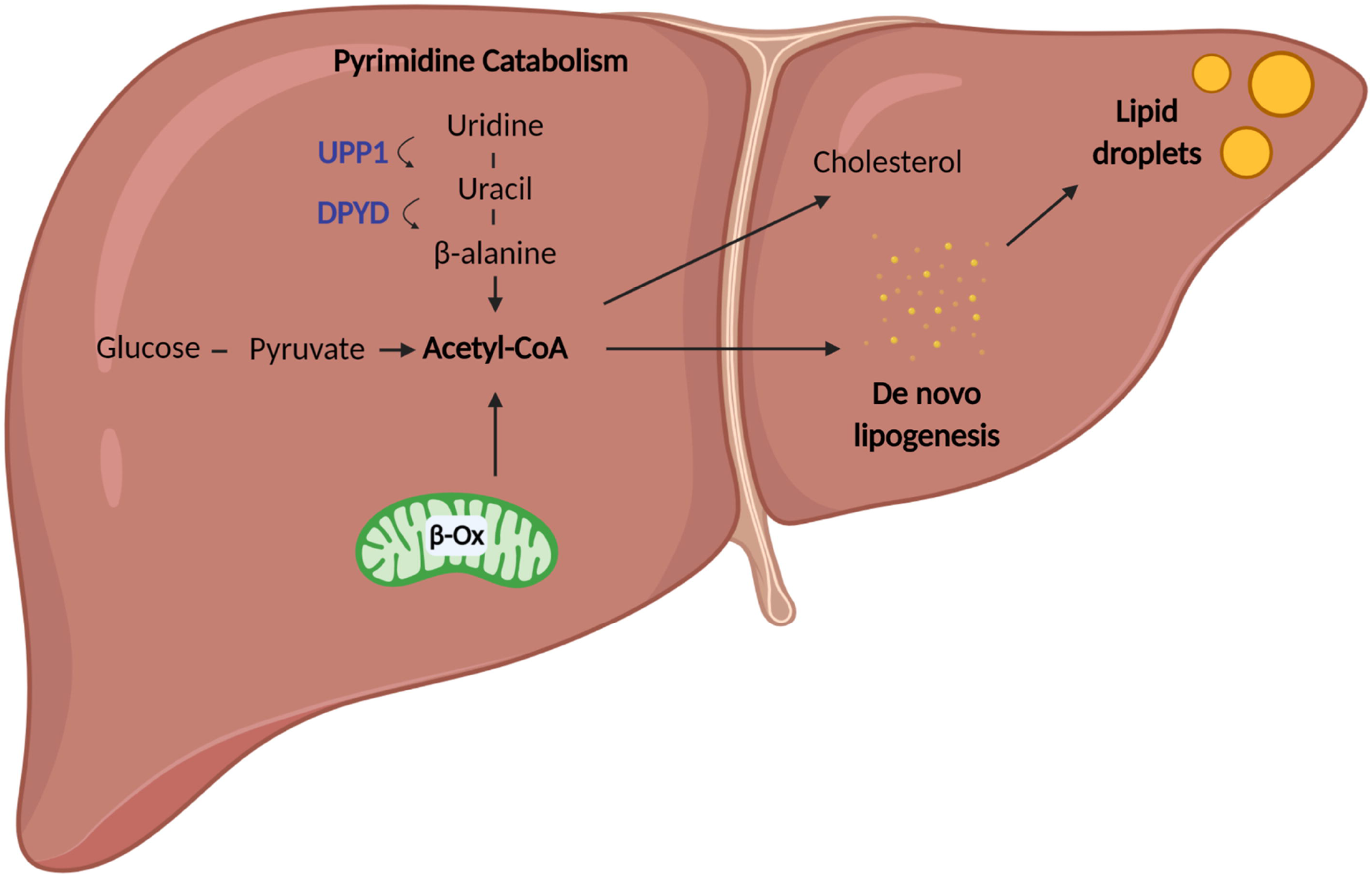
Characterization of human in vitro model of steatosis. A. Primary human hepatocytes treated with 200 µM palmitic and oleic acid demonstrate a significant increase in neutral lipid content as demonstrated by LipidTox stain (green). Scale bar represents 100 µM. B. Quantification of LipidTox Sum intensity demonstrates a significant correlation with free fatty acid concentration (R2=0.9469) C. RNAseq enrichment analysis highlights key pathways disturbed following free fatty acid treatment. Fatty acid metabolic process is the most significantly impacted pathway following free fatty acid treatment. D. A volcano plot describes the most significantly up (green) and down regulated (red) genes following exposure to palmitic and oleic acid. Dashed horizontal line indicates adjusted p-value of 0.05 while the dashed vertical lines represent 2 fold up and down regulation. ABCG1 is the most significantly upregulated gene and SLC25A47 is the most significant down regulated gene. E. Mitochondrial stress test on the seahorse instrument highlights an increase in oxygen consumption rate in the presence of 200 µM fatty acids. Mitochondrial stress test involves the addition of oligomycin A (2.0 µM) indicated at time interval A, addition of FCCP (2.0µM) at time interval B and addition of Rotenone Antimycin A (0.5µM) at time interval C.

In addition, markers of pathways associated with adipogenesis, fatty acid biosynthesis, unfolded protein response and production of NOS and ROS in macrophages were significantly influenced by the exposure to FFAs. Finally, markers of beta-alanine metabolism were markedly altered following FFA exposure (Figure 1C). *UPB1*, a gene encoding for the enzyme beta-ureidopropionase, was significantly downregulated following exposure to FFAs (adjusted p-value 0.025117). Beta-ureidopropionase is responsible for the final step of pyrimidine catabolism through its conversion of N-carbamyl-beta-alanine to beta alanine. The differential expression of this gene in our model system highlights the significance of pyrimidine catabolism in hepatocyte metabolic homeostasis.

### Functional characterization of DPYD inhibition in an in vitro primary human hepatocyte model of steatosis

With a characterized, *in vitro* model of steatosis, we next assessed how pharmacological inhibition of DPYD influenced the accumulation of lipids in cells. PHHs demonstrated reduced neutral lipid accumulation in a dose-dependent manner following Gimeracil exposure in the presence of FFAs; an IC50 value of 247.8 nM (Figure 2A and 2B). This difference in lipid burden due to DPYD inhibition was not an artifact of either cell density (Figure 2C) or viability (Figure 2D). The ability of Gimeracil to prevent hepatocyte lipid burden was observed across three different lines of PHHs (data not shown), indicating this was not an artifact of a particular donor line, but a true biological phenomenon.

**Figure 2.**
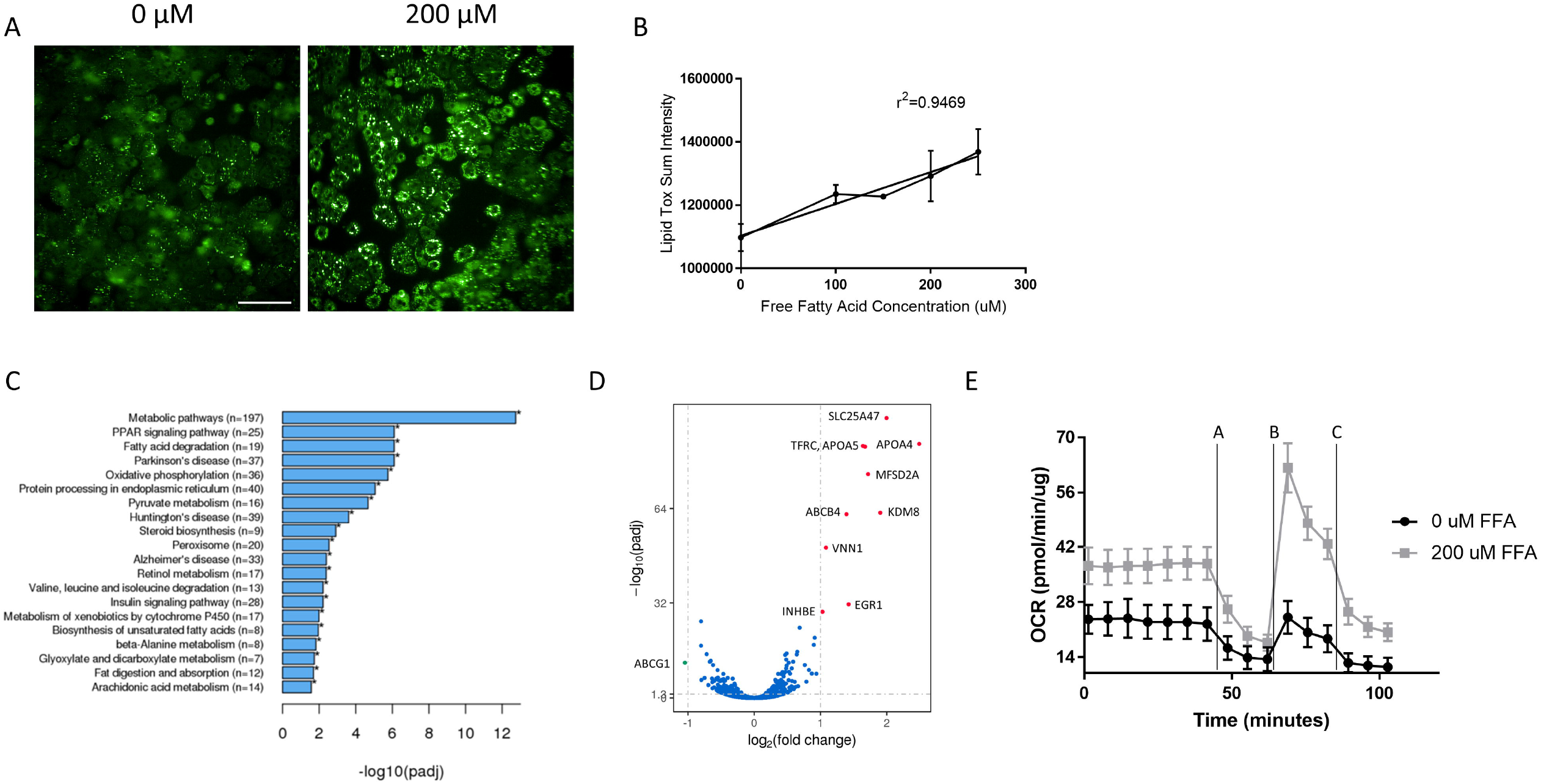
DPYD inhibition reduces lipid burden in an in vitro model of steatosis. A. While an increase in LipidTox signal intensity is observed in the presence of 200 µM free fatty acids, this effect is diminished in the presence of 250 nM Gimeracil, a DPYD inhibitor. Scale bar represents 100 µM. B. Quantification of lipid droplet accumulation in the hepatocytes with an Operetta Harmony image analysis pipeline (spot area per cell area) highlights a more than 2-fold increase in lipid droplet area with exposure to free fatty acids (square) relative to vehicle treatment (circle). Gimeracil exposure significantly reduces lipid droplet accumulation in both the free fatty acid and vehicle control conditons in a dose responsive fashion. The degree to which Gimeracil inhibits lipid accumulation in the presence of fatty acids is represented by an IC50 value of 247.8 nM with an R2 of 0.8481. C. Cell viability is not significantly impacted by either free fatty acid exposure (200 µM) or DPYD inhibitor up to a concentration of 500 nM as measured by a calcein AM stain confluence measurement quantified by the Incucyte Zoom analysis software. D. Neither 200 µM free fatty acid exposure or DPYD inhibition in the range of 50-500 nM is cytotoxic as measured by Ethidium homodimer-1 staining and quantification of confluence with the Incucyte Zoom analysis software.

The seahorse assay demonstrated that, in the presence of fatty acids, DPYD inhibition increased basal oxygen consumption with a maximal difference observed at 200 nm (adjusted p-value 0.0017) (Figure 3A). At this dose level of Gimeracil, the mitochondrial stress test revealed increases in maximal respiration (adjusted p-value <0.0001), spare respiratory capacity (adjusted p-value =0.0008) and ATP production (adjusted p-value =0.0002) (Figure 3C) relative to 200 µM fatty acid exposure in the absence of Gimeracil. Thus, these results suggest perturbation of the pyrimidine catabolism pathway enhances mitochondrial functionality and fatty acid oxidation.

**Figure 3.**
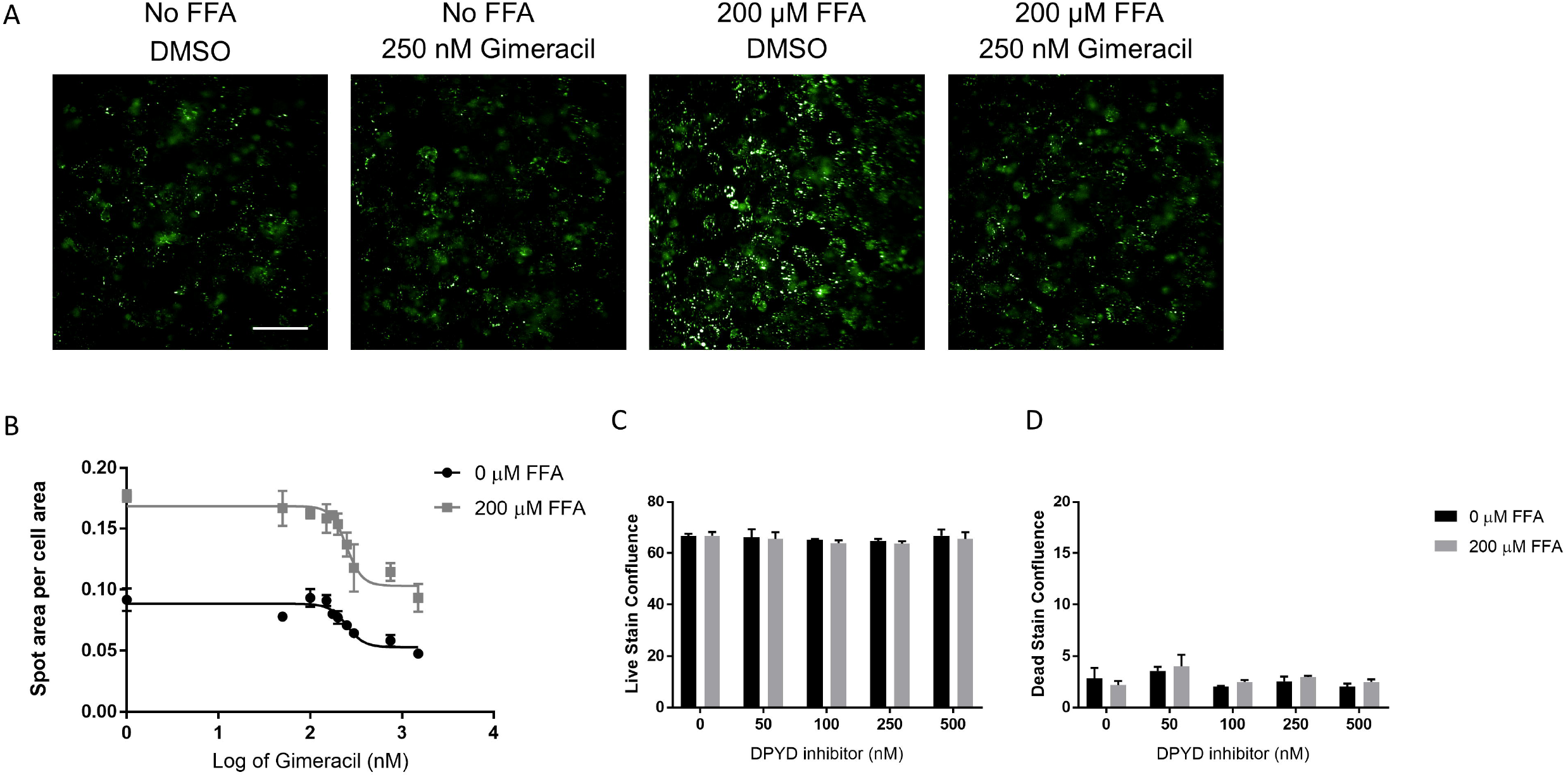
Extracellular flux analysis demonstrates enhanced mitochondrial functionality in primary human hepatocytes treated with the DPYD inhibitor Gimeracil. Mitochondrial stress test involves the addition of oligomycin A (2.0 µM) indicated at time interval A, addition of FCCP (2.0µM) at time interval B and addition of Rotenone Antimycin A (0.5µM) at time interval C. * indicates adjusted p-value<0.05 and ** indicates adjusted p-value<0.0001. A. Primary human hepatocytes treated with increasing concentrations of the DPYD inhibitor Gimeracil in the presence of 200 µM palmitic and oleic acid demonstrate a significant increase in oxygen consumption rate (OCR) at 50 nM (adjusted p-value 0.0173), 200 nM (adjusted p-value 0.0017) and 400 nM (adjusted p-value 0.0754). The increased observed at 1000 nM is not statistically significant (adjusted p-value 0.0754). B. A mitochondrial stress test was performed with 7 basal measurements collected every 7 minutes, followed by injections of 2 µM oligomycin, 2 µM FCCP and 0.5 µM Rotenone and Antimycin A. After injection, three measurements were recorded, each 7 minutes apart. Oxygen consumption rate (OCR) was determined following normalization to total cellular protein through a BCA assay. While FFA treatment (light gray square) increased OCR, the presence of 200 nM Gimeracil (dark gray triangle) further increased this rate of oxygen consumption relative to no treatment (black circle). C. Basal oxygen consumption increased with both FFA treatment (light gray, adjusted p-value 0.0064) and FFA and Gimeracil treatment (dark gray, adjusted p-value <0.0001). Maximal respiration, spare respiratory capacity and ATP production increased with FFA treatment (adjusted p-value <0.0001 for all three metrics) and increased further with Gimeracil treatment (adjusted p-value <0.0001 for all three metrics).

RNAseq analysis revealed minimal changes to the transcriptome as a result of DPYD inhibition, but those genes with differential transcript expression levels are key regulators of lipid biosynthesis and metabolism, lipid storage, glucose neogenesis and cholesterol synthesis. For example, the only differentially expressed transcripts, following exposure to 400 nM Gimeracil treatment in the presence of free fatty acids, were *HSD17B13, G0S2, MID1IP1, HMGCS1, CYP2B7P1, C19orf80, SLC25A47*, and *MALAT1* (Figure 4A). Several of these mRNA transcript changes demonstrated dose-dependent reductions following Gimeracil exposure in both the presence and absence of FFAs.

**Figure 4.**
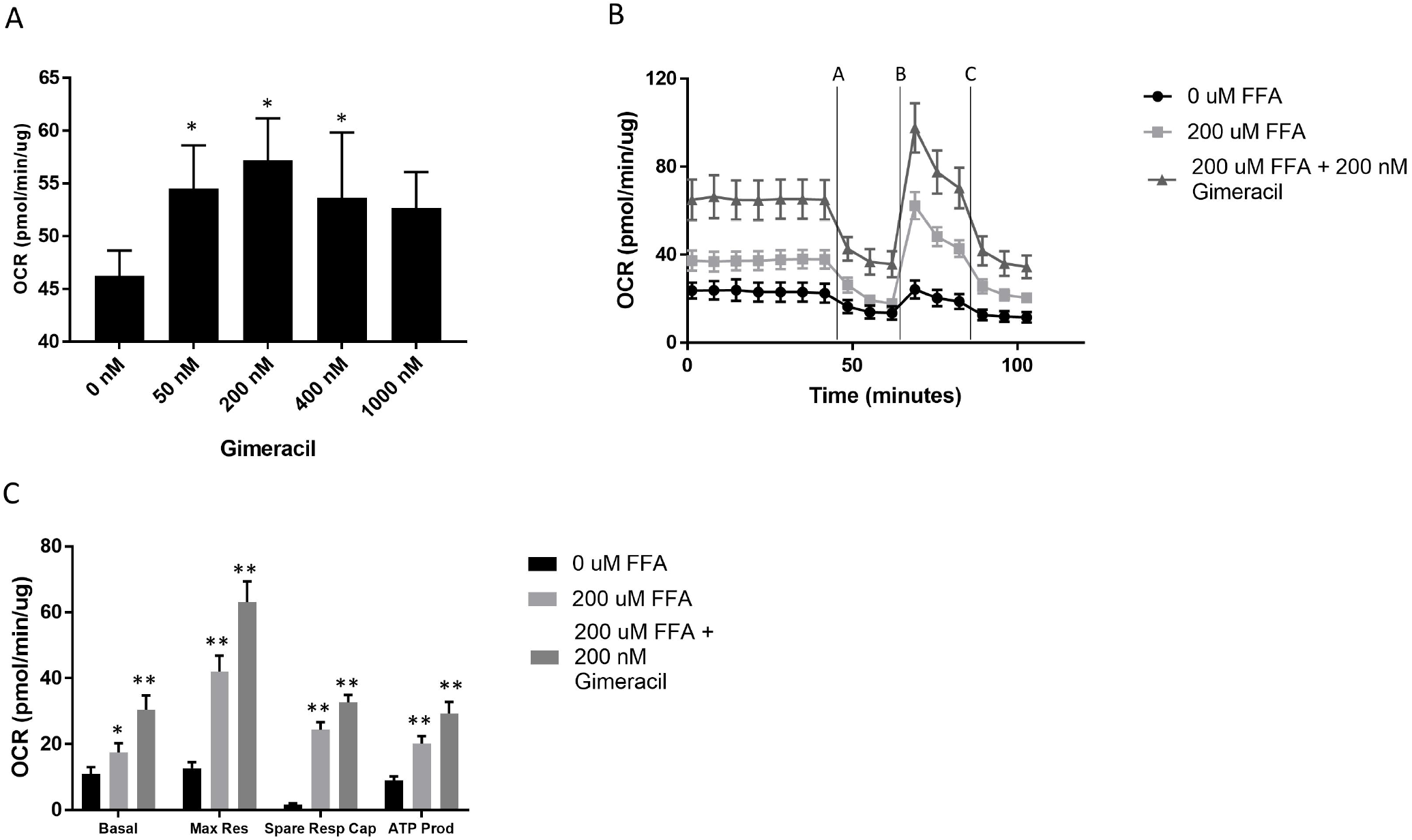
Gimeracil inhibition of DPYD perturbs metabolic processes related to cholesterol and lipid biosynthesis. A. A volcano plot describing the key genes with differential expression levels in the presence of 400 nM Gimeracil and 200 µM free fatty acids relative to Vehicle control with 200 µM free fatty acids. Log2 fold change and -log10 adjusted p values were plotted for the detected genes. HSD17B13 is the most differentially regulated gene, followed by TXNIP, G0S2, and MID1IP1. B. Pathway analysis reveals significant disruption in pathways associated with lipid, cholesterol and coenzyme metabolic processes in cells treated with 400 nM Gimeracil and 200 µM free fatty acids relative to vehicle control and 200 µM free fatty acids. N=3 samples per condition.

*Fatty acid synthase* (*FASN*) mRNA expression significantly decreased following Gimeracil exposure in both the presence and absence of FFAs (Table 1), with the most significant decrease observed for the highest dose of Gimeracil, (adjusted p-value 0.0123 and <0.0001 for vehicle and FFA exposure, respectively). While *FASN* mRNA expression decreased following FFA exposure in the absence of Gimeracil, the change was not significant (adjusted p-value 0.0607; Table 1). The *G0/G1 Switch Gene 2* (*G0S2*), a master regulator of hepatic lipid content through its inhibition of lipolysis (18), mRNA expression increased following exposure to FFAs (adjusted p-value=0.0016, Table 1) and decreased following exposure to Gimeracil (at the highest dose of 400 nM) in both the presence and absence of FFAs (adjusted p-value of 0.0123 and 0.0095, respectively; Table 1). *3-Hydroxy-3-Methylglutaryl-CoA Synthase 1* (*HMGCS1*), a rate-controlling enzyme in cholesterol biosynthesis, mRNA expression increased following exposure to FFA (adjusted p-value of 0.005; Table 1) and decreased in the presence of FFA at both the 200 and 400 nM dose of Gimeracil (adjusted p-value of 0.0001 and <0.0001, respectively; Table 1). The *hepatic lipid droplet protein hydroxysteroid 17-beta dehydrogenase 13* (*HSD17B13*) mRNA expression was significantly down-regulated at all dose levels of Gimeracil, in both the presence (adjusted p-values ≤0.0212 for all three dose groups; Table 1) and absence (adjusted p-values <0.001 for all three dose groups; Table 1), of FFA treatment. MID1 interacting protein 1 (MID1IP1) regulates hepatocyte lipogenesis through its regulation of Acetyl-CoA carboxylase 1 and *MID1IP1* mRNA was significantly down regulated with FFA exposure (adjusted p-value of 0.0005; Table 1). Expression levels of *MID1IP1* mRNA were also reduced with DPYD inhibition (Table 1), with a more dramatic dose responsive effect observed in the absence of FFA treatment (adjusted p-values of 0.0048 and p<0.0001 for 200 and 400 nM Gimeracil exposure, respectively). However, in the presence of FFAs, *MID1IP1* mRNA expression also significantly decreased at the highest dose level of Gimeracil (adjusted p-value 0.0114; Table 1). Also, significantly down-regulated with DPYD inhibition, in both the presence and absence of FFA treatment (adjusted p-values ≤0.0066 and ≤0.0177, respectively), was *Patatin Like Phospholipase Domain Containing 3* (*PNPLA3*); PNPLA3 is another lipid droplet associated protein. Finally, *thioredoxin interacting protein 1 (TXNIP*) mRNA expression, a critical regulator of hepatic glucose production (19), was significantly reduced as a function of Gimeracil treatment in the presence and absence of FFAs (adjusted p-values ≤0.0017 and 0.0101, respectively). Taken together, these results demonstrate the significant effect of Gimeracil on altering the metabolic profile of hepatocytes at the transcript level.

**Table 1.**
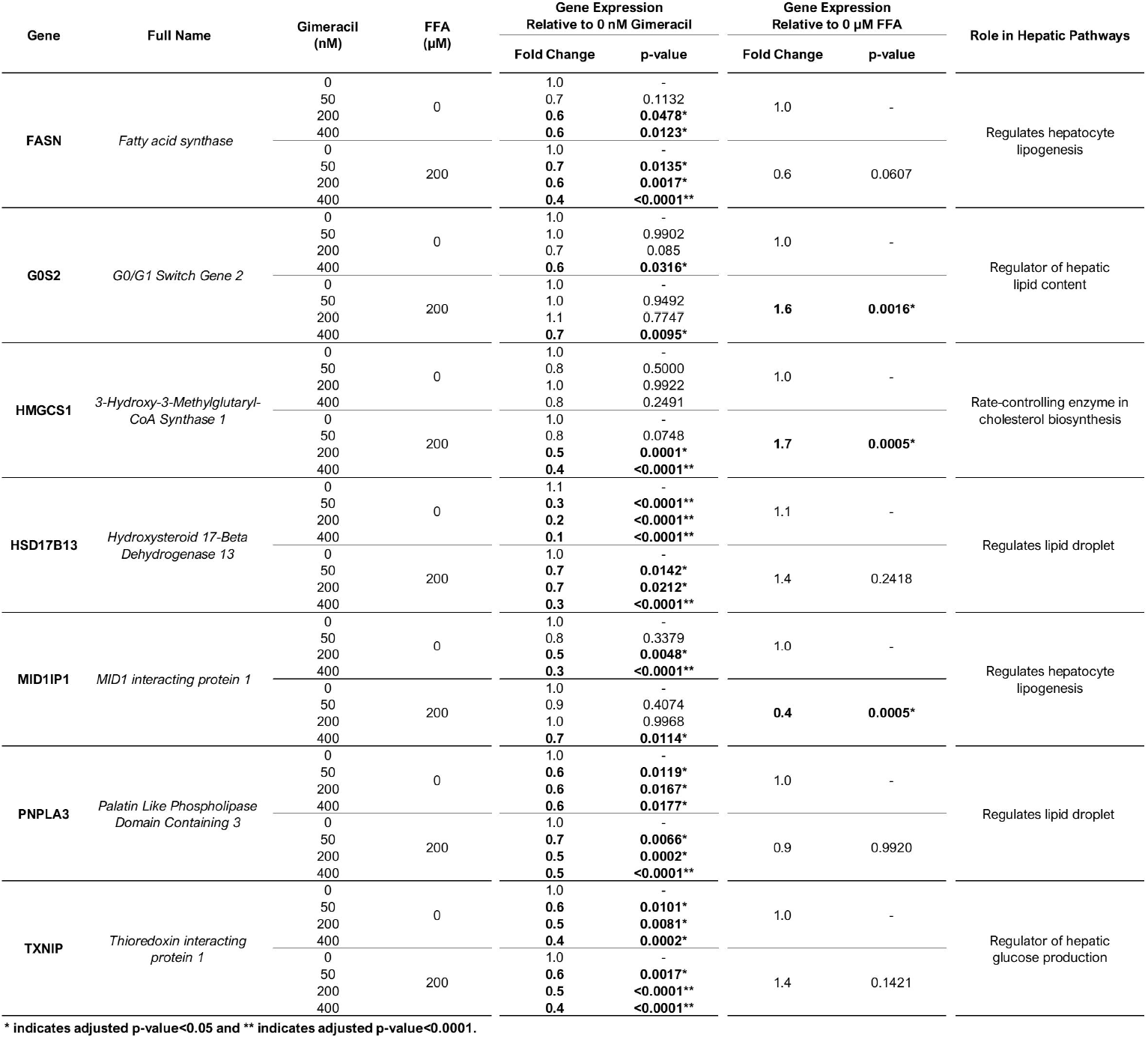
Validation of RNAseq differentially expressed genes through quantitative PCR. Fold change expression levels were calculated via delta delta Ct method relative to a.) 0 nM Gimeracil either in the presence or absence of FFA or b.) 200 µM FFA treatment relative to 0 µM FFA in the absence of Gimeracil. Significant differences in fold change expression level are denoted in bold with * indicating adjusted p value <0.05 and ** indicating p<0.0001. The role of each gene in hepatic metabolism is summarized in the right column of the table.

To help characterize the functions of DPYD variants, DPYD mutant forms were generated by site directed mutagenesis. As protein degradation was observed for both wild type and L310S mutant forms of DPYD, the concentration of the active protein was calculated based on the intensity of the ∼130 kDa band as a fraction of the total intensity of all bands on SDS-PAGE. The active protein concentrations were used for the determination of kinetic parameters of DPYD. The kinetic measurements of DPYD were carried out by varying concentrations of uracil. The obtained data fit best with a substrate inhibition model, yielding *k*_cat_ values of 65 ± 4 and 99 ± 5 min^-1^ for wild type and L310S mutant forms of DPYD (Figure 5B).

**Figure 5.**
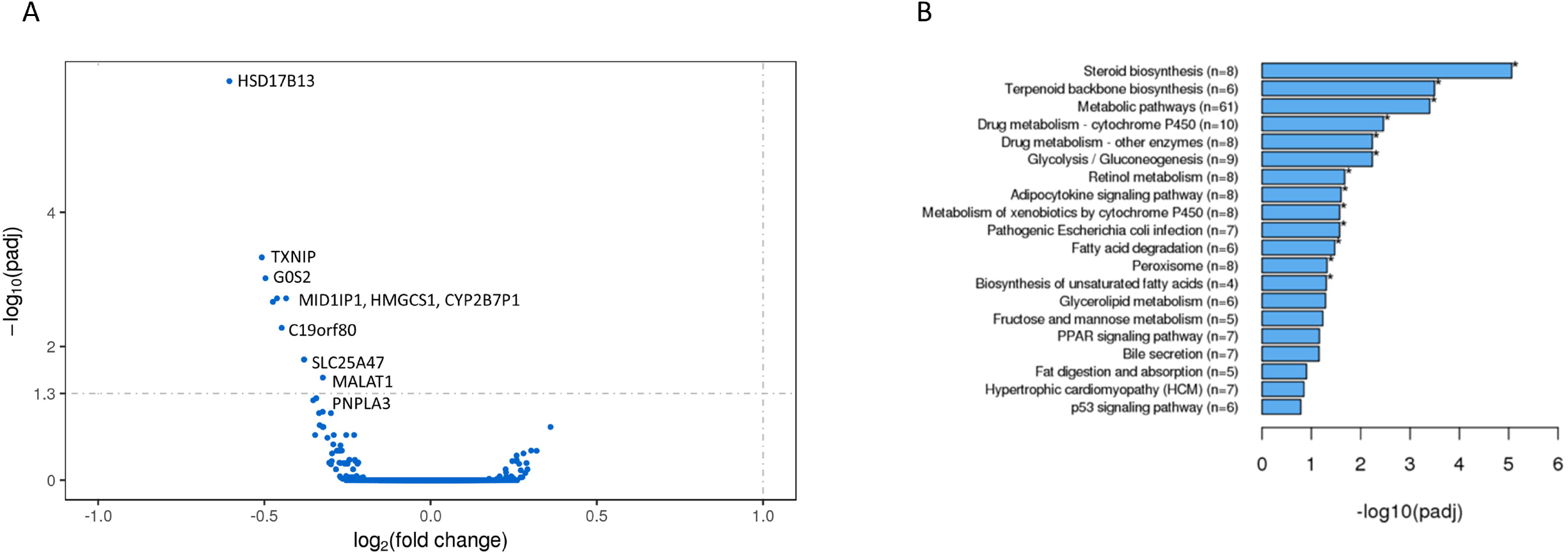
Purification and the activity assay of the recombinant DPYD. A. The visualization of purified wild type and L310S mutant forms of DPYD by SDS-PAGE. Lanes 1 & 4, molecular ladder (Thermo Scientific PageRuler Plus Prestained Protein Ladder); Lane 2, wild type DPYD; Lane 3, L310S mutant form of DPYD. B. Steady state kinetic measurement of wild type (●) and L310S mutant (▼) forms of DPYD. Enzymatic activities were measured in 100 mM potassium phosphate, pH 7.0, at 50 µM NADPH. The curves were fits of data to the steady state substrate inhibition (19).

Screening of hepatocarcinoma cell lines based on their expression levels of both *UPP1* and *DPYD* mRNA transcript demonstrated that the SNU449 cell line had the optimal mRNA expression levels of both enzymes as measured by their relative ratio of 1, whereas PHHs demonstrated a ratio of 1.1 (Figure 6A). This cell line responded to FFA exposure with an increase in lipid content and Gimeracil treatment significantly reduced lipid accumulation in a dose-responsive manner as well (R^2^=0.8560 and IC50 of 19.74 nM) (Figure 6B and 6C). Next-Generation sequencing revealed successful generation of a homozygous L310S DPYD mutant line and a homozygous, PAM-edit only, control line through homology directed repair (Figure 7). In addition, employing NHEJ methodology, a heterozygous knock-out mutant line was generated with allele A possessing a D308_F309delinsV mutation and allele B possessing a S306_K307_delinsL* mutation. PROVEAN predicted allele A would be functionally deleterious with a score of -17.289 (20) and the premature stop codon on allele B yields a nonsense mutation (Figure 7B). Western blot analysis revealed almost total ablation of DPYD protein expression in the KO mutant line (Figure 7C).

**Figure 6.**
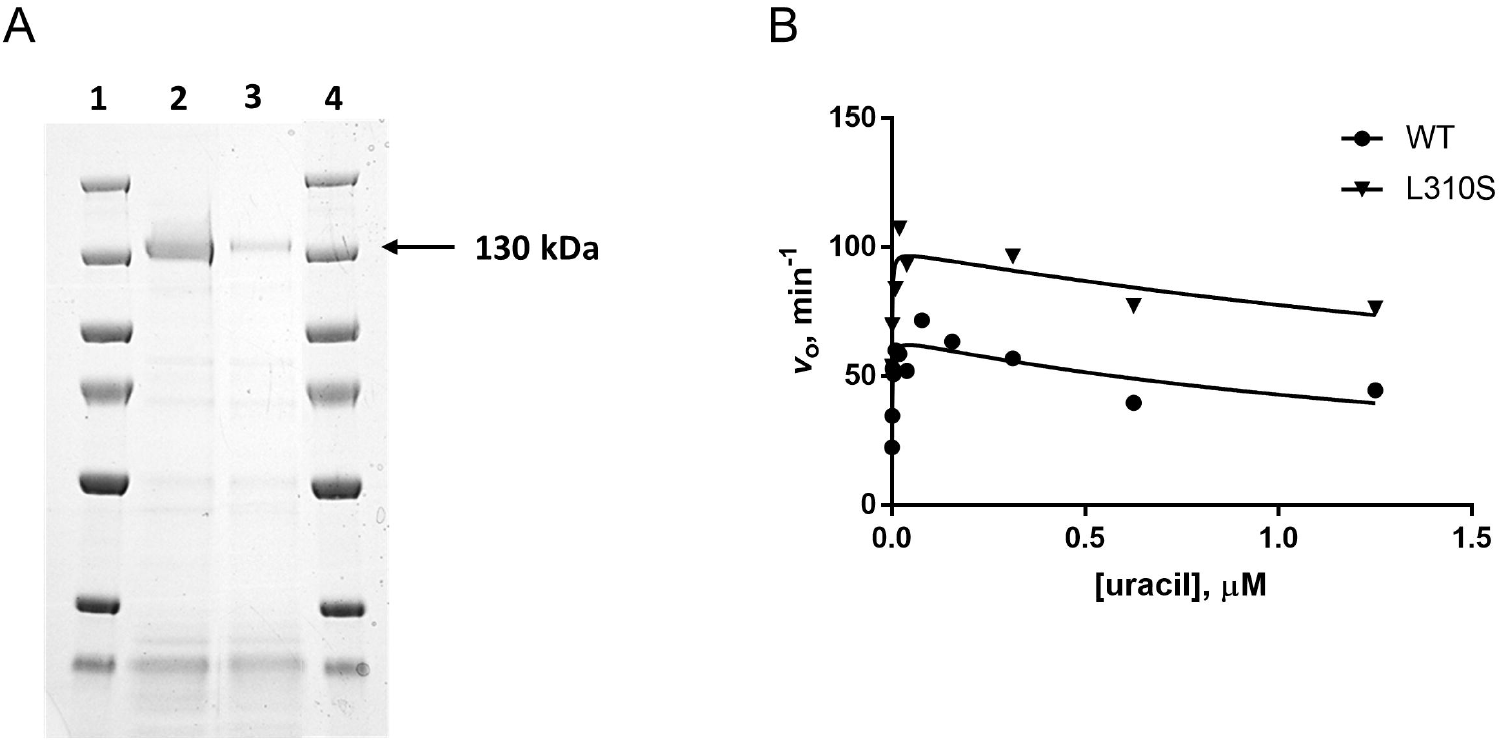
Selection of SNU449 cell line for modeling DPYD gain and loss of function mutations. A. Digital droplet PCR was performed to assess the relative expression levels of UPP1 and DPYD normalized as a percentage of the geometric mean of GAPDH, UBC and RSP20. The ratio of their normalized expression was calculated and compared to primary human hepatocytes (PHH). The relative ratio of UPP1:DPYD in PHH (BioIVT lot WWQ) was 1.1, which was most closely represented by the SNU449 line with a ratio of 1.0. B. LipidTox signal intensity increased with 100 µM free fatty acid exposure. DPYD inhibition decreased the LipidTox spot area per cell area in a dose responsive manner (IC50 of 19.74 nM and R2 of 0.865) suggesting DPYD inhibition is protective against free fatty acid-induced neutral lipid accumulation in SNU449 cell line. C. Representative images of SNU449 cell line treated with fatty acids in the presence and absence of 100 nM DPYD inhibitor demonstrate a significant increase in lipid accumulation (represented by the LipidTox stain) with FFA treatment that is further reduced in the presence of 100 nM DPYD inhibitor. Scale bar represents 100 µM.

**Figure 7.**
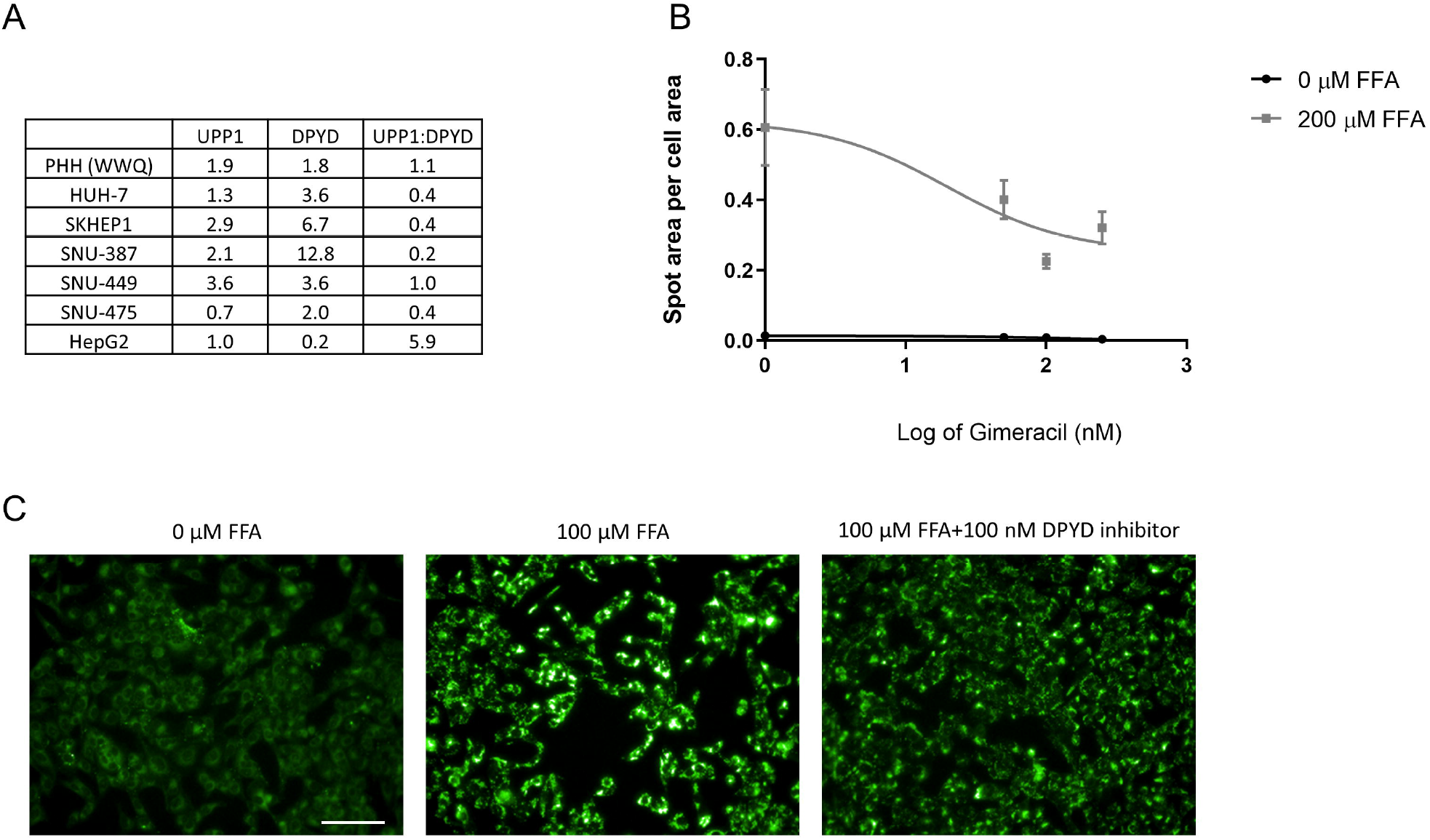
CRISPR-Cas9 engineered SNU449 mutant line characterization. A. Genotype of three mutant SNU449 lines; Knock-out (KO) mutant line generated through NHEJ and L310S line and synonymous PAM edited control line both generated through homology-directed repair. Site of PAM sequence edit, site of non-synonymous HDR edit for L310S generation and Guide RNA sequence highlighted for reference. B Miseq targeted amplicon genotyping reveal homozygosity of L310S mutant and control line with 99.62 and 99.40% sequence identity, respectively, with reference sequence described in tile A. KO mutant line demonstrates heterozygosity with 46.45% sequence identity with allele A (D308_F309delinsV) and 52.17% sequence identity with allele B (S306_K307_delinsL*). C. Western blotting describes loss-of DPYD protein expression in KO line. L310S line demonstrates reduced DPYD protein expression relative to control. Equal protein loading demonstrated through β-actin expression.

The L310S mutant line also appeared to have reduced expression relative to the control line. Functional characterization of the three lines revealed, relative to the control, a significant increase in lipid burden in the L310S mutant line at baseline (adjusted p-value 0.0130) and a significant reduction in lipid level in the KO line (adjusted p-value 0.0213) (Figure 8). Following exposure to FFAs, all three cell lines exhibited an increase in lipid burden, with the L310S demonstrating the highest LipidTox signal intensity as measured by spot area per cell area (with an adjusted p-value <0.0001 relative to control line treated with fatty acids) (Figure 8).

**Figure 8.**
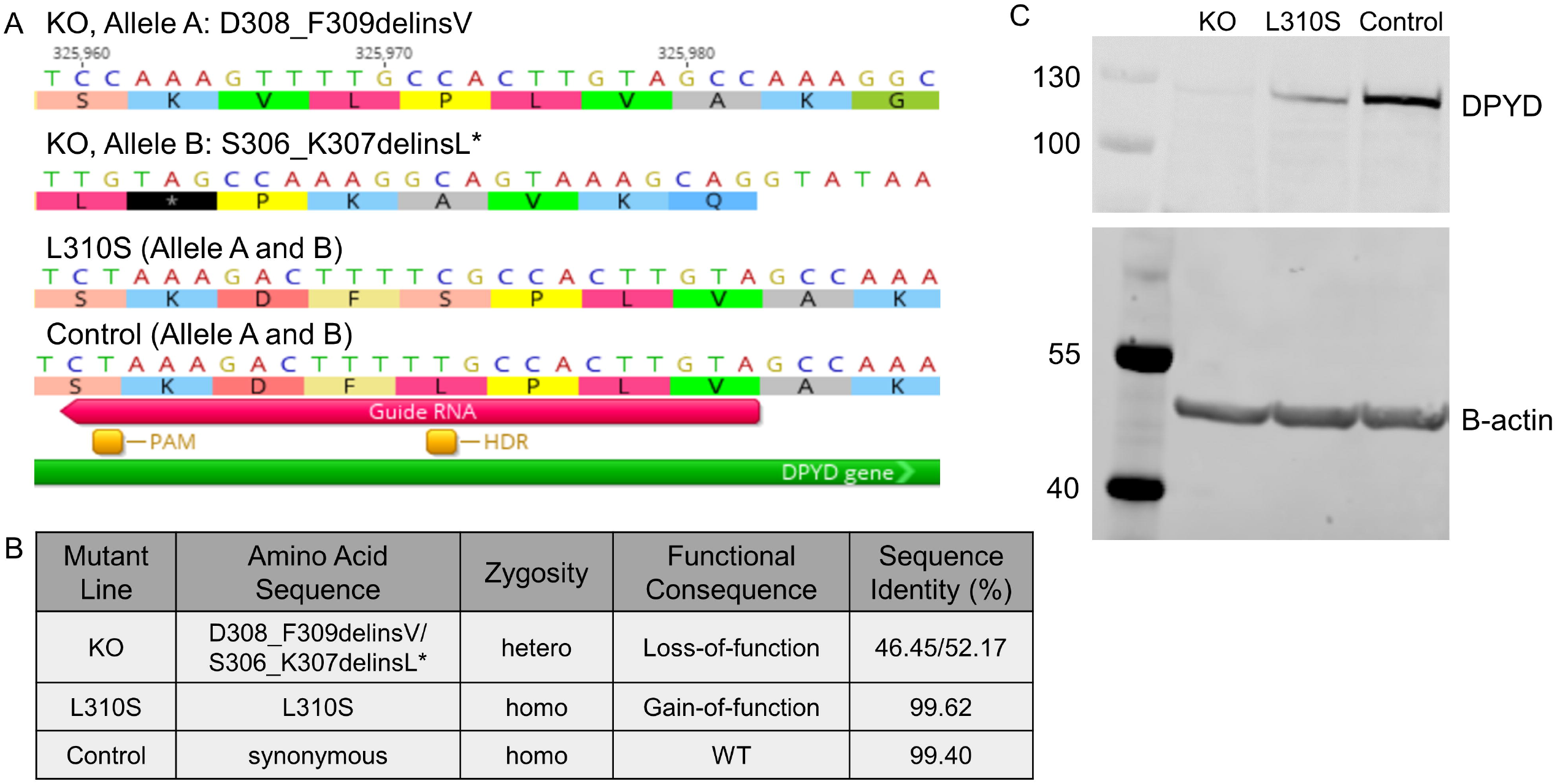
Mutant SNU449 phenotypic characterization. A. LipidTox neutral lipid stain reveals that under basal conditions, mutant SNU449 CRSIPR-Cas9 engineered cell lines show altered lipid burdens. The L310S line demonstrates an increased lipid burden and the KO line shows a reduced lipid burden. Under 25 µM free fatty acid treatment, all three lines demonstrate an increased lipid burden, with the L310S line demonstrating the greatest signal intensity for LipidTox neutral lipid stain. Scale bar represents 100 µM in all images, except for inlays where the scale bar represents 50 µM. Image thresholding was optimized for both vehicle and fatty acid conditions to highlight differences across lines. Within a treatment condition, image thresholding is constant. B. Quantitative image analysis demonstrates a significant increase in LipidTox spot area per cell area in the L310S line relative to control (adjusted p-value of 0.0130). The KO line shows a significantly reduced lipid burden under basal conditions as measured by a significant decrease in neutral lipid spot area per cell area relative to control (adjusted p-value of 0.0213). Upon free fatty acid exposure the L310S line demonstrates a significant increase in lipid accumulation as measured by LipidTox spot area per cell area (adjusted p-value <0.0001).

Mitochondrial function was compared across these mutant lines (Figure 9). Extracellular flux analysis revealed a significant reduction in the rate of glycolysis in the KO line (adjusted p-value<0.0001) (Figure 9A) and a significantly lower rate of oxygen consumption in the L310S line (adjusted p-value 0.0083) (Figure 9B). A mitochondrial stress test revealed an increase in the spare respiratory capacity in the KO line (adjusted p-value 0.0377) (Figure 9C) and a decrease in maximal respiration rate in the L310S line (adjusted p-value 0.0201) (Figure 9D). These results highlight that, relative to the control cell line, the loss-of-function line had a positive change in energy utilization and improved mitochondrial capacity, whereas the gain-of-function mutation showed impaired mitochondrial function, likely due, impart, to increased lipid burden.

**Figure 9.**
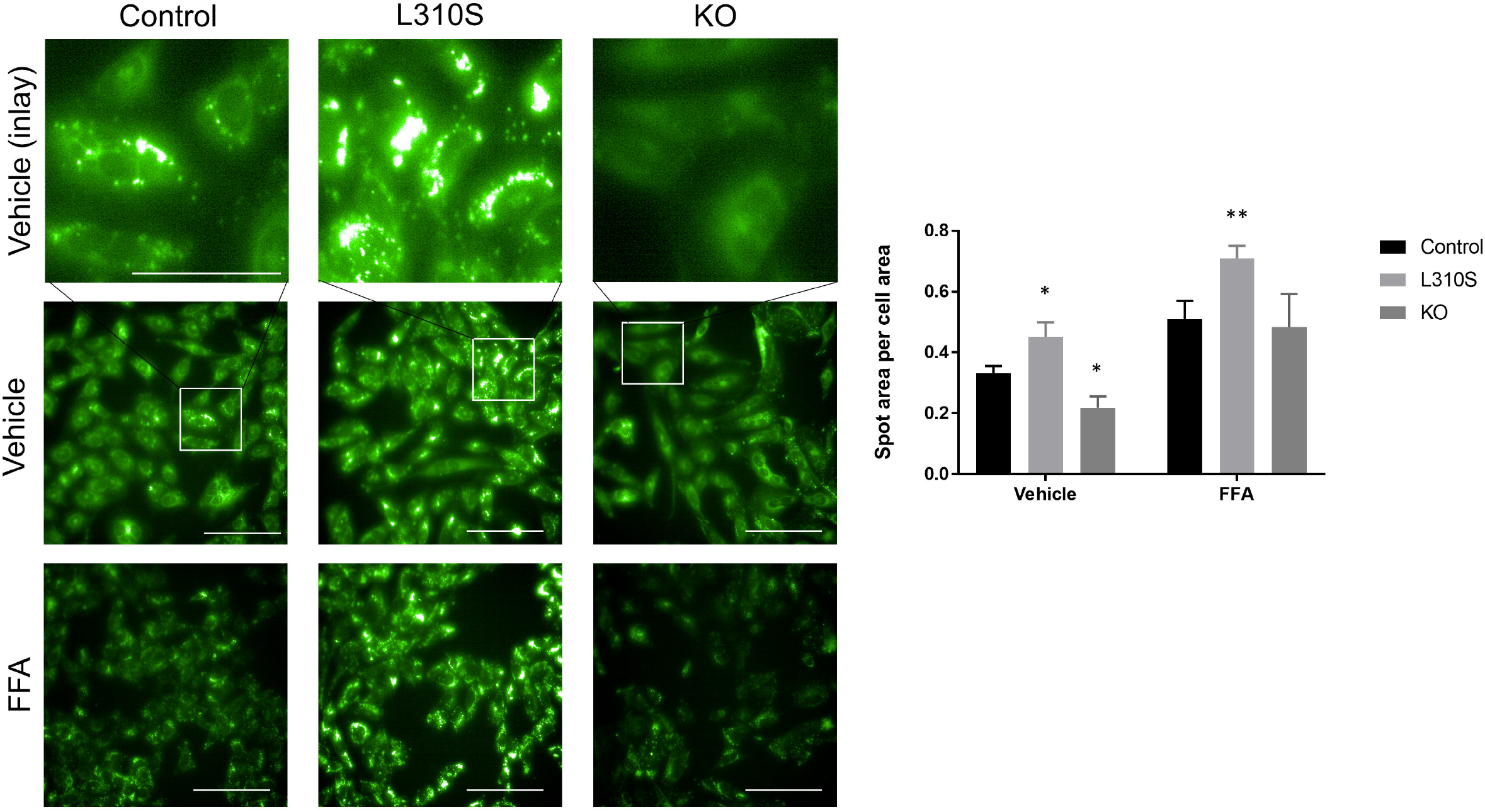
Evaluation of mitochondrial functionality in the CRISPR-Cas9 engineered cell lines through extracellular flux analysis with a mitochondrial stress test involving the addition of oligomycin A (2.5 µM) indicated at time interval A, addition of FCCP (2.0µM) at time interval B and addition of Rotenone Antimycin A (0.5µM) at time interval C. Black (circle) indicates control, light gray (square) indicates L310S line and dark gray (triangle) indicates KO line. * indicates p-value < 0.05. A. Extracellular acidification rate (mpH/min/µg) (ECAR) is a measurement of glycolysis in the mitochondrial stress test. The KO loss of function DPYD line demonstrates a significant reduction in the rate of glycolysis relative to the control and L310S line at all measured time intervals (adjusted p-value ≤0.0065). B. Oxygen consumption rate (pmol/min/µg) (OCR) is a measure of mitochondrial respiration and L310S demonstrates a reduced rate of oxygen consumption at baseline and no increase in oxygen consumption with the addition of FCCP. The KO line demonstrates similar rates of oxygen consumption and responsiveness to too compounds, with the exception of the addition of FCCP. The KO line responds more rapidly to FCCP addition with a more significant increase in OCR. C. There is no significant difference in coupling efficiency (Coupling Eff) across the three lines, but the KO line demonstrates a more significant increase in spare respiratory capacity (Spare Resp Cap) relative to the control line (adjusted p-value 0.0377). D. The L310S line has a reduced basal OCR (adjusted p-value 0.0083) and a reduced rate of maximal respiration (Max Resp) (adjusted p-value 0.0201) relative to control.

We then assessed whether the gain and loss-of-function mutant SNU449 lines had differential expression of genes involved in lipogenesis, lipolysis, and gluconeogenesis, which had been previously implicated in the PHH cultures treated with Gimeracil (Figure 10). Relative to the control line, *FASN* mRNA expression was significantly reduced in the KO line (adjusted p-value 0.0044) under basal conditions, but this difference was not maintained in the presence of FFAs (Supplemental figure 1). *G0S2* mRNA expression was significantly increased in both the L310S and KO mutant lines, in both the presence and absence of FFAs (adjusted p-values <0.0001). There was no change in the mRNA expression of *G0S2* in the control line following FFA treatment. In the presence of FFA, *HSD17B13* mRNA expression was significantly increased in the KO line (adjusted p-values ≤0.0150); no significant change was detected in the control line. While *PNPLA3* mRNA expression also did not change significantly in the control line following FFA treatment, *PNPLA3* expression was significantly reduced in the KO line in the absence of FFAs (adjusted p-value 0.0093) and significantly increased in the L310S line in the presence of FFAs (adjusted p-value 0.0101). *TXNIP* mRNA expression was significantly upregulated following FFA treatment in the control line (adjusted p-value <0.0001), and even further increased in the KO line relative to the control under both basal conditions as well as following exposure to FFA (adjusted p-values <0.0001). Additionally, relative to the control line, the L310S gain-of-function line also demonstrated an increase in *TXNIP* mRNA expression under basal conditions (adjusted p-value 0.0261). Again, compared to the control line, *UPP1* mRNA expression was significantly increased in the KO line under both basal conditions as well as following FFA exposure (adjusted p-values ≤0.0244). These results highlight how the metabolic phenotype of the hepatocarcinoma cell line SNU449 are altered as a function of genetic perturbation of DPYD.

**Figure 10.**
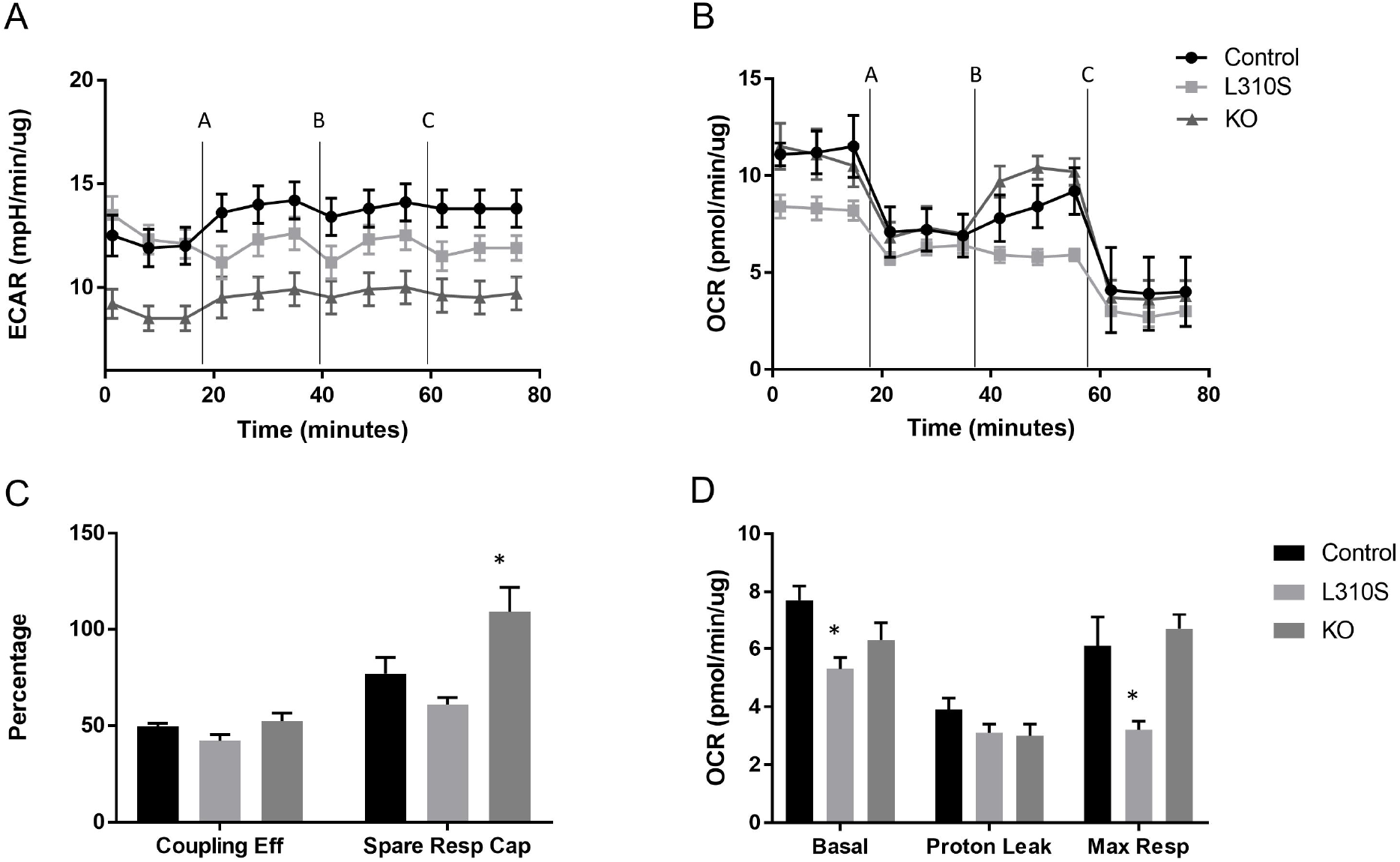
qPCR demonstrates differences in gene expression across three DPYD CRISPR-cas9 engineered cell lines; control (gray), control with FFA (black), L310S (pink), L310S with FFA (red), KO (light blue) and KO with FFA (dark blue). Data is normalized to control with or without FFA depending on treatment group. Under basal conditions, in the absence of free fatty acid exposure, DPYD, HMGCS1, HSD17B13, and MID1IP1 expression is consistent across the three cell lines. FASN expression is significantly downregulated in the L310S line relative to control (adjusted p-value 0.0044). G0S2 is significantly upregulated in both the KO and L310S line relative to control (adjusted p-value <0.0001 for both lines). PNPLA3 expression is significantly decreased in the KO line (adjusted p-value 0.0093). TXNIP expression is significantly increased in the L310S line (adjusted p-value 0.0261) and the KO line relative to control (adjusted p-value <0.0001). UPP1 expression is significantly increased in the KO line relative to control (adjusted p-value 0.0244). Following exposure to free fatty acids, DPYD, FASN, HMGCS1, and MID1IP1 expression remains consistent across lines. G0S2 expression significantly increased in both the L310S and the KO line relative to control with FFA (adjusted p-value <0.0001). HSD17B13 expression significantly increases in the KO line relative to control with FFA (adjusted p-value of 0.0105). PNPLA3 expression significantly increases in the L310S line relative to control with FFA (adjusted p-value of 0.0101). TXNIP expression is significantly increased in the KO line relative to control with FFA (adjusted p-value <0.0001). UPP1 expression is significantly increased in the KO line relative to control with FFA(adjusted p-value <0.0001).

## Discussion

The pharmacologic and genetic perturbation of DPYD enzymatic activity described herein reveal its role in regulating a hepatocyte’s metabolic phenotype *in vitro*. DPYD inhibition ameliorated lipid accumulation in an *in vitro* model of diet-induced steatosis in a dose dependent manner; the observed difference in lipid phenotypes corresponded with an enhanced rate of mitochondrial respiration and decreased expression of genes associated with de novo lipogenesis, lipolysis, lipid storage, and gluconeogenesis. A gain-of-function mutation in DPYD in a hepatocarcinoma cell line resulted in increased lipid burden, impaired mitochondrial respiration, and increased mRNA expression of several genes associated with lipid storage. A loss-of-function mutation in DPYD resulted in a decreased rate of glycolysis and increased mRNA expression of genes associated with gluconeogenesis and decreased mRNA expression of genes associated with lipogenesis and lipid storage. As no human genetic association between DPYD and NASH has so far been identified, it is possible that compensatory mechanisms maintain hepatic energy homeostasis in the context of mutant DPYD enzymatic activity. However, it is possible that pharmacologic perturbation of DPYD activity may be a promising therapeutic strategy for the amelioration of steatosis and the prevention of NASH.

As the rate limiting enzyme in uracil catabolism, DPYD regulates the metabolism of uracil into beta-alanine, which is further metabolized into a key energy source of the cell, Acetyl CoA. Acetyl CoA is the primary substrate of the Krebs cycle wherein its carbon atoms are oxidized for energy production (21). Other sources of Acetyl CoA include β-oxidation of fatty acids and oxidative decarboxylation of pyruvate following glycolysis (22). The abundance of Acetyl CoA not only reflects the energy state of the cell, but also regulates the acetylation profiles of various proteins including histones, which influence energy metabolism, autophagy and mitosis through direct and epigenetic gene regulation (23). In the presence of the DPYD inhibitor, Gimeracil, the hepatic abundance of Acetyl CoA was presumably decreased, representing a “fasted” state in which fatty acids are oxidized to replenish mitochondrial Acetyl CoA levels. The increase in fatty acid oxidation was reflected by the increased rate of mitochondrial respiration measured through extracellular flux analysis (Figure 3B and 3C), the reduction of neutral lipids in the cytosol (Figure 2A and 2B), and decreased mRNA expression of *G0S2* (Table 1), a gene encoding a protein which inhibits lipolysis (18). With a decreased lipid burden, mRNA expression of genes encoding proteins associated with lipid metabolism, such as *HSD17B13* and *PNPLA3*, are significantly down regulated (Table 1). In addition, lipogenesis, or the synthesis of fatty acids from Acetyl CoA, is reduced with DPYD inhibition as demonstrated by the decreased mRNA expression of *FASN* and *MID1IP1* (Table 1). DPYD inhibition also decreased the mRNA expression of genes encoding enzymes required for glucose production such as, *TXNIP*; possibly a result of reduced stores of Acetyl CoA. Finally, DPYD inhibition decreased the mRNA expression of *HMGCS1*, a gene encoding a rate limiting enzyme in the synthesis of cholesterol from Acetyl CoA (24), further providing supporting evidence that DPYD regulates energy stores of the cell likely in an Acetyl-CoA-dependent manner. Together, these *in vitro* results highlight the critical role DPYD plays in hepatic metabolic homeostasis, through its regulation of a central metabolite, Acetyl CoA, in its catabolism of uracil.

While DPYD mediated catabolism of uracil impacts cellular Acetyl CoA levels, it also directly influences plasma uridine levels. Uridine administration improves glucose tolerance in aging and diet-induced animal models of insulin resistance (13), while also preventing lipid accumulation in drug-induced cell models of hepatic steatosis (9). It has been demonstrated that adipocyte lipolysis, impacted by Xbps1 overexpression, is dependent on uridine synthesis, suggesting that driving adipocyte-mediated synthesis of uridine could be a promising treatment option for the mitigation of obesity (25). While the mechanism by which uridine production triggers systemic weight loss in mice is still unknown, it is understood that plasma uridine levels influence thermoregulation during fasting and feeding cycles (13), and that a uridine-driven decrease in body temperature correlates with greater lipid oxidation and oxygen consumption (26). Therefore, the protective effect of DPYD inhibition in an *in vitro* model of diet-induced steatosis is likely two-fold; (i) by increasing hepatic uridine levels, DPYD inhibition promotes glucose tolerance and lipolysis and (ii) by decreasing a major metabolite of the cell, Acetyl CoA, DPYD inhibition promotes utilization of existing energy stores.

Given the human genetic variability that exists in DPYD (exemplified by the toxicity to the chemotherapeutic 5-FU), it is important to consider whether gain-of and loss-of-function mutations in DPYD regulate differences in the metabolic phenotype of hepatocytes. *In vitro* cell culture models provide a controlled environment to interrogate the proximal effects of DPYD mutations in a specific cellular population while minimizing secondary, compensatory mechanisms present in whole tissues and organ systems. We identified a gain-of-function mutation in DPYD that drives an increased lipid burden in a hepatocarcinoma cell line; this altered phenotype coincided with mitochondrial dysfunction and increased expression of genes associated with lipid storage. In contrast, a loss-of-function mutation in DPYD demonstrated a reduced lipid burden under basal conditions with corresponding reduced rates of glycolysis and increased expression of genes associated with gluconeogenesis. Interestingly, both mutations drive not only a significant increase in the expression of *G0S2* mRNA, which thereby inhibits lipolysis under fasting conditions, but also an increased response to endoplasmic reticulum (ER) stress (27). Therefore, it is possible that the increased level of *G0S2* mRNA expression is impacted by differential mechanisms in the two DPYD mutant lines. One possible explanation may be that the significant accumulation of lipids in the gain-of-function mutant DPYD line give rise to an increase in reactive oxygen species and ER stress, leading to increased G0S2 expression as a compensatory mechanism to minimize cellular stress through the prevention of lipolysis. However, in the loss-of-function DPYD line, there is a reduction in energy stores in the cell as demonstrated by decreased glycolysis and increased expression of mRNA for genes involved in glucose production. Increased G0S2 expression in the loss-of-function line may inhibit lipolysis in this “fasting” state to conserve energy stores (28).

While the loss-of-function DPYD mutant line demonstrated reduced lipid burden under basal conditions, this phenotypic difference was not maintained in the presence of free fatty acids, as lipid accumulation was not significantly reduced relative to the control line. This observation contradicts the pharmacological experiments in the wild-type SNU449 line wherein DPYD inhibition reduced lipid accumulation following FFA exposure. Therefore, it is possible that the loss-of-function mutation is not protective against FFA induced lipid accumulation due to compensatory mechanisms which regulate energy homeostasis in the absence of DPYD enzymatic activity. For example, under basal conditions, the loss-of-function line demonstrates a phenotype consistent with energy conservation through its reduced rate of glycolysis, inhibition of lipolysis and induction of gluconeogenesis. The addition of a new energy source in the form of FFAs may not initiate a rapid shift in the cells’ metabolic profile from a state of energy conservation to consumption. Future work assessing the metabolic phenotype of the loss-of-function line over a time course longer than 24 hours as well as varying concentrations of fatty acid exposure may reveal new insights into the functional consequence of a DPYD loss of function mutation.

As the conclusions drawn from this body of work were based on data collected from human *in vitro* cell model systems, a future effort should be dedicated to assessing the functional effects of DPYD inhibition in a rodent model of hepatic steatosis. Due to both the a.) biological diversity between humans and rodents, as well as b.) the increased complexity of the *in vivo* setting, it is possible that DPYD-inhibition *in vivo* may not have a functional consequence on hepatic lipid accumulation. However, a body of preclinical work demonstrates uridine levels influence hepatic lipid accumulation, providing scientific rationale for DPYD inhibition as a significant driver of liver lipid levels and a potential clinical target for mitigating hepatic steatosis. Our application of human cell model systems overcomes the limitations associated with previous preclinical models by demonstrating pyrimidine catabolism influences human hepatic steatosis *in vitro*. Thus, employing both pharmacologic and genetic perturbation methods in both human cell lines and human primary cells, we identified a novel role for DPYD in human hepatic lipid homeostasis. Since the DPYD inhibitor, Gimeracil, is already deemed safe and tolerable in the clinic when delivered in combination with the chemotherapeutic 5-FU for the treatment of various cancers, Gimeracil may be a promising therapeutic for the alleviation of hepatic steatosis.

## Materials and Methods

### Functional characterization of DPYD inhibition in an in vitro primary human hepatocyte model of steatosis

Collagen I coated plates were used for all cell culture experiments described herein. Primary human hepatocytes (PHHs, BioIVT, lot WWQ) were overlaid with 0.25mg/mL Matrigel. 24 hours after seeding, cells were treated with free fatty acids (FFAs) consisting of a 1:1 ratio of BSA-conjugated Oleic acid and Palmitic acid (100, 150, 200 and 250 µM) with concurrent treatment with Gimeracil (50, 100, 150, 175, 200, 250, 300, 750, 1500 nM). 48 hours after seeding, cells were fixed, stained with LipidTox (ThermoFisher, H34475), imaged on a Perkin-Elmer Operetta and analyzed with a Harmony image analysis pipeline identifying nuclei count, LipidTox sum intensity/cell and LipidTox spot area/cell area. Experiments were repeated with additional lines of PHHs (BioIVT, lots JEL and RVQ). Additional samples were assessed for viability with a Live/Dead assay (ThermoFisher, L3224) 24 hours after treatment with 200 µM FFAs and DPYD inhibitor bromovinyl uracil (50, 100, 250 and 500 nM). Cells were imaged and analyzed on an Incucyte Zoom.

For the mitochondrial respiration assays, the Agilent Seahorse XFe96 Analyzer was used with the Mito Stress Test assay according to the manufacturer’s recommended protocol. PHHs were seeded onto XF96 cell culture microplates. 24 hours after seeding, cells were treated with Gimeracil (50, 200, 400 and 1,000 nM). 24 hours later, cell media was changed to Krebs-Henseleit buffer (pH7.4) containing 2.5 mM glucose, 0.5 mM carnitine and 5 mM HEPES. 30 µL of Palmitate-BSA or BSA control was added to the microplates immediately prior to loading onto the instrument. All data was normalized to protein content with a BCA assay.

For RNAseq experiments, PHHs were treated with Gimeracil (50, 200, and 400 nM) for 24 hours and then treated with FFAs. 24 hours after FFA exposure, RNA was isolated from cells with the Qiagen RNeasy kit according to manufacturer’s protocol. Sequencing was performed by Novogene and sequencing reads were aligned to human transcriptome reference (GRCh38.p12) and gene counts were quantified via STAR after quality control on fastq files. The differential expression analysis was conducted with DESeq2 R package and pairwise comparisons were made between groups. The significant criteria are padj < 0.05 and |log2(FoldChange)| > 1. The differentially expressed genes were then applied to clusterProfiler for Gene Ontology (GO) enrichment analysis with default settings. GO terms with padj < 0.05 were considered significantly enriched by differential expressed genes. In addition, clusterProfiler was also used to test the statistical enrichment of differential expression genes in KEGG pathways. KEGG terms with padj < 0.05 are considered as significant enrichment (18).

Quantitative PCR was performed to validate RNA Sequencing results (Taqman primers in Supplemental Table 3). Following reverse transcription with Superscript Vilo IV kit, RT-PCR was performed with QuantStudio 5 instrument. The geometric mean of three housekeeping genes (GAPDH, β-actin and UBC) was used to normalize the data. The delta delta Ct method was used to calculate fold change in expression relative to DMSO treatment (0 nM Gimeracil) with and without 200 µM FFA exposure. BSA was used as the vehicle control for FFA treatment.

#### In vitro *functional characterization of human DPYD mutants*

Two gene block fragments coding for DPYD (Supplemental Table 1) were inserted into a pET28a(+) expression vector using the In-Fusion HD Cloning kit. Site directed mutagenesis (Agilent’s QuickChange II kit) was performed using mutagenesis primers detailed in Supplemental Table 1. DHα5 cells were transformed with mutant expression vectors and confirmed through Sanger sequencing at Genewiz (Cambridge, MA).

WT and mutant vectors were used to transform BL21(DEC3)pLysS cells in Luria-Bertani broth medium containing kanamycin (50 µg/ml). After the overnight growth, the starter culture was used to inoculate 200 ml of culture medium containing 50 µg/ml kanamycin, 100 µM uracil, 10 µM of sodium sulfide, 10 µM ammonium ferric citrate, 10 µM of flavin adenine dinucleotide, and 10 µM flavin mononucleotide. When the culture reached an optical density at 600 nm, (8 to 10 hours at 30 °C), 0.5M final IPTG was added. Cells were harvested and stored at – 80 °C.

Purification steps were carried out at 4 °C. The cell paste was suspended in BugBuster solution (Millipore 70921) that was buffered in 25 mM Tris-Cl, containing 250 mM NaCl, 0.2 mg/ml lysozyme, 0.1 mM phenylmethylsulfonyl fluoride, 0.05 mg/ml Benzonase I, 1.25 mM DTT and 0.2% Tween 20, pH 8.0 (buffer A). The resuspended cell pellet was incubated on ice and underwent ultra-centrifugation for 15 minutes at 40,000g. The supernatant was collected and loaded onto a PD 10 column (pre-equilibrated with buffer A) that was prepacked with 2 ml slurry cobalt beads following the manufacture’s instruction and then eluted with 10 volumes buffer A, followed by another 10 volumes of the same buffer with additional 0.5 M NaCl. Subsequently, another 10 volumes of buffer A with an additional 10 mM imidazole was used to elute the column. Finally, 2 volumes of buffer A with 0.3 mM imidazole, followed by another 2 volumes of buffer A with 0.5 mM imidazole was applied. The fractions of the imidazole elution were pooled together and concentrated using an Amicon ultra centrifugal filters (MWCO = 15,000 Da). After removal of the precipitated protein by centrifugation, concentration was assessed via Nanodrop.

Initial velocities of the DPYD reaction were assayed fluorometrically by monitoring the depletion of NADPH at *E*_x_/*E*_m_ = 340/460 nM at 25 °C in a 384-well plate using SpectroMax. Continuous measurement was carried out for 30 minutes. The extinction coefficient of NADPH was determined by the *E*_x_/*E*_m_ = 340/460 read out of NADPH, which was serially diluted from 50 µM to 50 nM. The standard reaction contains 100 mM potassium phosphate at pH 7.0, 0.5 mM dithiothreitol (DTT), 0.1 mM NADPH, varying concentrations of uracil (i.e., 16 nM to 20 µM), and 2-6 nM enzyme, in a total volume of 12 µL. The kinetic data was fit into a Michaelis-Menten or a substrate inhibition model (19).

### Screening of hepatocarcinoma cell lines

Hepatocarcinoma cell lines were selected for their gene expression levels of DPYD and UPP1 enzymes: HUH7 (JCRB0403), Skhep1 (HTB-52), SNU387 (CRL-2237), SNU449 (CRL-2234), and SNU475 (CRL-2236) (all purchased from the ATCC, except HUH7 purchased from Xenotech). RNA was extracted from cell pellets with the Qiagen RNeasy mini isolation kit, reverse transcribed with the VILO IV reverse transcription kit and quantified gene expression levels of GAPDH, B-actin, RSP-20, UPP1 and DPYD through digital droplet PCR (ddPCR) (Supplemental Table 3).

Cell lines were seeded in RPMI media containing 10% FBS and changed to serum-free RPMI after 24 hours. 48 hours after seeding, cells were treated with FFAs and bromovinyl uracil (50, 100 ad 250 nM). Exposure to FFAs above 100 µM was cytotoxic. Plates were fixed and stained with LipidTox and imaged following the same protocol as outlined for PHHs.

### Generation of gain and loss-of-function DPYD mutant SNU449 lines with CRISPR-Cas9 homology directed repair and non-homologous end joining

Two guide RNAs were designed to target the DPYD L310S locus (20). These guide RNAs were generated with the Precision GuideRNA synthesis kit with primers ordered from IDT (Supplemental Table 1). Single strand DNA oligos were designed with phosphoorothioate linkages to promote intracellular stability (Supplemental Table 1). Guide RNA quality and quantity was determined with both a Small RNA bioanalyzer kit and Broad Range RNA Qubit kit. Optimal concentrations of guide RNA, Cas9 enzyme and ssDNA oligo (10 µg of guide RNA, 5 µg of Cas9 enzyme and 10 µg of ssDNA oligo) were combined to maximize both non-homologous end joining (NHEJ) and homology directed repair (HDR) efficiency. Guide RNA, Cas9 enzyme and ssDNA oligo were combined in SF solution and incubated on ice for 20 minutes. Cas9 enzyme and the control Nucleofector GFP plasmid were incubated together in SF solution as a control.

The SNU449 cell line was pelleted (200,000 cells per reaction) and resuspended in SF solution containing guide RNA, Cas9 enzyme and ssDNA oligo. Nucleofection (96 well Nucleofector plate, 20uL) was performed according to the manufacturer’s suggested protocol for HepG2 cells. Following nucleofection, cells were resuspended in 10% FBS in RPMI media. Half of the cells were genotyped and the remaining were expanded. Genomic DNA was extracted with directPCR lysis buffer and Proteinase K (0.4mg/mL). Proteinase K was inactivated at 95°C and DNA was genotyped with a ddPCR-based assay (primers/probes outlined in Supplemental Table 1). Thermocycling conditions were performed according to the manufacturer’s instructions for the probe supermix containing no dUTP. Guide 1 was selected for future experiments as NHEJ editing efficiencies were superior at 98% efficiency.

HDR efficiencies for the generation of the L310S mutation were estimated at 9%. The bulk population was then expanded for one passage, seeded at single cell density and expanded for 3 weeks in culture. Approximately 30% of seeded wells propagated and underwent ddPCR genotyping. Only cultures which demonstrated a homozygous L310S mutation, PAM edit only mutation and homozygous NHEJ mutations were expanded up to a T-75 flask and cryopreserved.

Next generation sequencing libraries were generated according to published protocol (21) and first and second round primers are detailed in Supplemental Table 2. Libraries were pooled together, diluted to 4 nM, and denatured in NaOH according to the Illumina Miseq protocol. Libraries were further diluted to 6.25 pM and spiked with 6 pM denatured PhiX at a ratio of 3:1. Samples were run with an Illumina Micro V2 kit with single read processing and generated over 300,000 reads per sample. NGS sequencing identified a homozygous L310S mutant line, a control, homozygous PAM edit only line and a heterozygous NHEJ line with an allele with a 19 base pair deletion and a premature stop codon (S306_K307_delinsL*) and an allele with a three base-pair deletion, D308_F309delinsV. The functional consequence of this mutation was assessed with PROVEAN analysis software (22).

Cell lysate was isolated with RIPA buffer containing protease inhibitor cocktail and quantified with a BCA assay. Protein lysate underwent electrophoresis (4-12% Bis-Tris NuPage gradient gel in MOPS buffer) and was transferred to a nitrocellulose membrane (NuPage Transfer buffer with 20% methanol). The blot was blocked in 5% Milk in tris-buffered saline with 0.1% Tween 20 (TBST) and then incubated in 5% Milk TBST containing mouse anti-DPYD (Sigma, WH0001806M1) and rabbit anti-β-actin (Cell Signaling, 4967S) primary antibodies. Blot was rinsed in TBST, incubated with donkey anti-mouse IgG cross-adsorbed DyLight 800 (ThermoFisher, SA5-10172), and goat anti-rabbit IgG cross-adsorbed DyLight 680 secondary antibodies (ThermoFisher, 35568), rinsed in TBST and imaged on a Li-Cor imager.

Mutant cell lines were thawed following genotyping with Illumina Miseq instrument and cultured for three passages. Cells were seeded in 10% FBS in RPMI. 24 hours after seeding, cell media was changed to serum free RPMI. For the seahorse assay, after 24 hours in serum free media, cells were analyzed with the mitochondrial seahorse assay as described for PHHs. For imaging and mRNA analysis, after 24 hours in serum free media, cells were treated with 25 µM FFAs. 24 hours after treatment, the 96 well plate was fixed, stained with LipidTox and imaged on the Operetta as described previously. RNA was isolated, reverse transcribed and mRNA levels quantified with Taqman primers via qPCR as described previously. Data was normalized to the geometric mean of three housekeeping genes (β-actin, GAPDH and HPRT1). The delta-delta ct method was used to calculate fold change relative to the control, PAM-edit only mutant line with or without free fatty acid exposure.

### Statistical Analysis

All data presented in figures represent mean +/- standard deviation. Imaging, seahorse mitochondrial assay and quantitative PCR data were analyzed with a two-way analysis of variance with an alpha of 0.05 and Tukey’s post-hoc testing for multiple comparisons. For dose response curves, a 3 parameter, non-linear fit model was used (log(inhibitor) vs response) (Prism). The default Benjamini-Hochberg method was used to control False Discovery Rate (FDR) of the DESeq2 identified deferential expressed genes.

## Abbreviations

5-FU: (5-Fluorouracil)
CRISPR-Cas9: (clustered regularly interspaced short palindromic repeats associated protein 9)
DPYD: (dihydropyrimidine dehydrogenase)
FASN: (fatty acid synthase)
FXR: (farsenoid X receptor)
G0S2: (G0/G1 Switch Gene 2)
HMGCS1: (3-Hydroxy-3-Methylglutaryl-CoA Synthase 1)
HSD17B13: (hepatic lipid droplet protein hydroxysteroid 17-beta dehydrogenase 13)
MID1IP1: (MID1 interacting protein 1)
NAFLD: (nonalcoholic fatty liver disease)
NASH: (non-alcoholic steatohepatitis)
PHH: (primary human hepatocyte)
PNPLA3: (Patatin Like Phospholipase Domain Containing 3)
TXNIP: (thioredoxin interacting protein 1)
UPP1: (uridine phosphorylase-1)

## Acknowledgement

The authors thank the Department of Translational Safety and Bioanalytical Sciences at Amgen for their support throughout the duration of this postdoctoral research. The authors would also like to thank Ingrid Rulifson (Department of the Cardiometabolic Disorders, Amgen Inc) for her insightful revisions.

## Figures

**Supplemental Figure 1.**
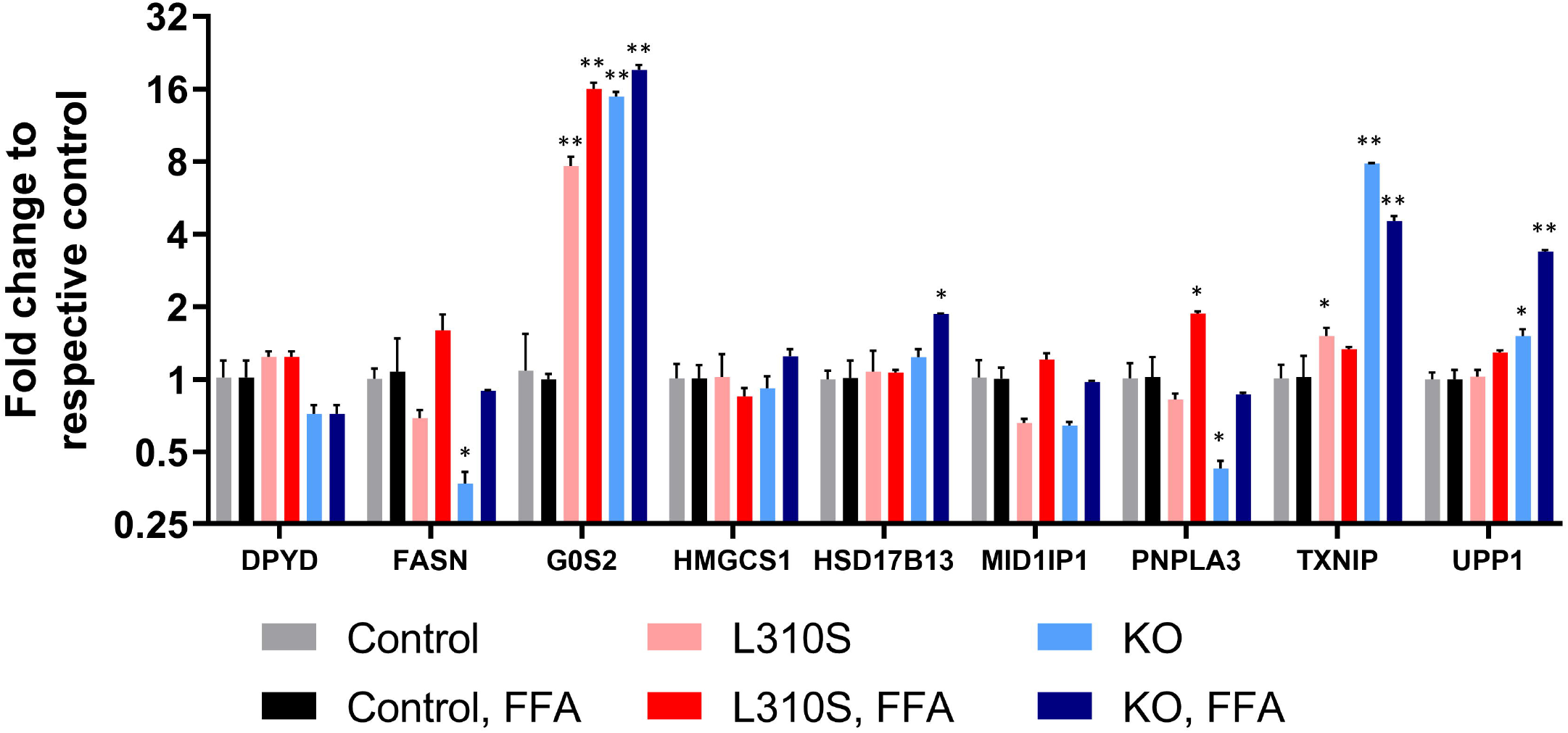
Gene expression changes in SNU449 control line exposed to free fatty acids relative to no treatment. There is no significant change in expression of DPYD, FASN, G0S2, HMGCS1, HSD17B13, MID1IP1, PNPLA3 or UPP1. There is a significant increase in expression of TXNIP following free fatty acid treatment relative to no treatment (adjusted p-value <0.0001).

Supplemental Table 1. The nucleic acid sequences of the two gene block fragments used in molecular cloning of DPYD.

## References

1. El-Zayadi A-R. Hepatic steatosis: a benign disease or a silent killer. World journal of gastroenterology. 2008;14(26):4120–6.

2. Oseini AM, Sanyal AJ. Therapies in non-alcoholic steatohepatitis (NASH). Liver international : official journal of the International Association for the Study of the Liver. 2017;37 Suppl 1(Suppl 1):97–103.

3. Younossi Z, Anstee QM, Marietti M, Hardy T, Henry L, Eslam M, George J, Bugianesi E. Global burden of NAFLD and NASH: trends, predictions, risk factors and prevention. Nature Reviews Gastroenterology & Hepatology. 2018;15(1):11–20.

4. Doycheva I, Issa D, Watt KD, Lopez R, Rifai G, Alkhouri N. Nonalcoholic Steatohepatitis is the Most Rapidly Increasing Indication for Liver Transplantation in Young Adults in the United States. J Clin Gastroenterol. 2018;52(4):339–46.

5. Nikolaou N, Gathercole LL, Marchand L, Althari S, Dempster NJ, Green CJ, van de Bunt M, McNeil C, Arvaniti A, Hughes BA, Sgromo B, Gillies RS, Marschall H-U, Penning TM, Ryan J, Arlt W, Hodson L, Tomlinson JW. AKR1D1 is a novel regulator of metabolic phenotype in human hepatocytes and is dysregulated in non-alcoholic fatty liver disease. Metabolism. 2019;99:67–80.

6. Le TT, Ziemba A, Urasaki Y, Hayes E, Brotman S, Pizzorno G. Disruption of uridine homeostasis links liver pyrimidine metabolism to lipid accumulation. Journal of lipid research. 2013;54(4):1044–57.

7. Khusial RD, Cioffi CE, Caltharp SA, Krasinskas AM, Alazraki A, Knight-Scott J, Cleeton R, Castillo-Leon E, Jones DP, Pierpont B, Caprio S, Santoro N, Akil A, Vos MB. Development of a Plasma Screening Panel for Pediatric Nonalcoholic Fatty Liver Disease Using Metabolomics. Hepatology Communications. 0(0).

8. Liu Y, Zhang Y, Yin J, Ruan Z, Wu X, Yin Y. Uridine dynamic administration affects circadian variations in lipid metabolisms in the liver of high-fat-diet-fed mice. Chronobiology International. 2019;36(9):1258–67.

9. Le TT, Urasaki Y, Pizzorno G. Uridine prevents tamoxifen-induced liver lipid droplet accumulation. BMC Pharmacol Toxicol. 2014;15:27.

10. Lebrecht D, Vargas-Infante YA, Setzer B, Kirschner J, Walker UA. Uridine supplementation antagonizes zalcitabine-induced microvesicular steatohepatitis in mice. Hepatology. 2007;45(1):72–9.

11. Le TT, Urasaki Y, Pizzorno G. Uridine prevents fenofibrate-induced fatty liver. PLoS One. 2014;9(1):e87179.

12. Urasaki Y, Pizzorno G, Le TT. Chronic Uridine Administration Induces Fatty Liver and Pre-Diabetic Conditions in Mice. PLOS ONE. 2016;11(1):e0146994.

13. Deng Y, Wang ZV, Gordillo R, An Y, Zhang C, Liang Q, Yoshino J, Cautivo KM, De Brabander J, Elmquist JK, Horton JD, Hill JA, Klein S, Scherer PE. An adipo-biliary-uridine axis that regulates energy homeostasis. Science. 2017;355(6330).

14. Wigle TJ, Tsvetkova EV, Welch SA, Kim RB. DPYD and Fluorouracil-Based Chemotherapy: Mini Review and Case Report. Pharmaceutics. 2019;11(5).

15. Peng L, Piekos S, Guo GL, Zhong X-b. Role of farnesoid X receptor in establishment of ontogeny of phase-I drug metabolizing enzyme genes in mouse liver. Acta Pharmaceutica Sinica B. 2016;6(5):453–9.

16. Maekawa K, Saeki M, Saito Y, Ozawa S, Kurose K, Kaniwa N, Kawamoto M, Kamatani N, Kato K, Hamaguchi T, Yamada Y, Shirao K, Shimada Y, Muto M, Doi T, Ohtsu A, Yoshida T, Matsumura Y, Saijo N, Sawada J. Genetic variations and haplotype structures of the DPYD gene encoding dihydropyrimidine dehydrogenase in Japanese and their ethnic differences. J Hum Genet. 2007;52(10):804–19.

17. Kobayakawa M, Kojima Y. Tegafur/gimeracil/oteracil (S-1) approved for the treatment of advanced gastric cancer in adults when given in combination with cisplatin: a review comparing it with other fluoropyrimidine-based therapies. Onco Targets Ther. 2011;4:193–201.

18. Ma Y, Zhang M, Yu H, Lu J, Cheng KKY, Zhou J, Chen H, Jia W. Activation of G0/G1 switch gene 2 by endoplasmic reticulum stress enhances hepatic steatosis. Metabolism. 2019;99:32–44.

19. Chutkow WA, Patwari P, Yoshioka J, Lee RT. Thioredoxin-interacting protein (Txnip) is a critical regulator of hepatic glucose production. J Biol Chem. 2008;283(4):2397–406.

20. Choi Y, Chan AP. PROVEAN web server: a tool to predict the functional effect of amino acid substitutions and indels. Bioinformatics (Oxford, England). 2015;31(16):2745–7.

21. Wilson K, Hess J, Zhang G, Brunengraber H, Tochtrop G. Metabolism of Beta-Alanine in Rat Liver: Degradation to Acetyl-CoA and Carboxylation to 2-(aminomethyl)-malonate. The FASEB Journal. 2017;31(1_supplement):655.3-.3.

22. Bhagavan NV, Ha C-E. Chapter 16 - Lipids I: Fatty Acids and Eicosanoids. In: Bhagavan NV, Ha C-E, editors. Essentials of Medical Biochemistry. San Diego: Academic Press; 2011. p. 191–207.

23. Pietrocola F, Galluzzi L, Bravo-San Pedro José M, Madeo F, Kroemer G. Acetyl Coenzyme A: A Central Metabolite and Second Messenger. Cell Metabolism. 2015;21(6):805–21.

24. Vock C, Doring F, Nitz I. Transcriptional regulation of HMG-CoA synthase and HMG-CoA reductase genes by human ACBP. Cell Physiol Biochem. 2008;22(5-6):515–24.

25. Deng Y, Wang ZV, Gordillo R, Zhu Y, Ali A, Zhang C, Wang X, Shao M, Zhang Z, Iyengar P, Gupta RK, Horton JD, Hill JA, Scherer PE. Adipocyte Xbp1s overexpression drives uridine production and reduces obesity. Molecular metabolism. 2018;11:1–17.

26. Peters GJ, van Groeningen CJ, Laurensse EJ, Lankelma J, Leyva A, Pinedo HM. Uridine-induced hypothermia in mice and rats in relation to plasma and tissue levels of uridine and its metabolites. Cancer Chemother Pharmacol. 1987;20(2):101–8.

27. Wang Y, Zhang Y, Qian H, Lu J, Zhang Z, Min X, Lang M, Yang H, Wang N, Zhang P. The g0/g1 switch gene 2 is an important regulator of hepatic triglyceride metabolism. PLoS One. 2013;8(8):e72315.

28. Heckmann BL, Zhang X, Xie X, Liu J. The G0/G1 switch gene 2 (G0S2): Regulating metabolism and beyond. Biochimica et Biophysica Acta (BBA) - Molecular and Cell Biology of Lipids. 2013;1831(2):276–81.

29. Hill CM, Waightm RD, Bardsley WG. Does any enzyme follow the Michaelis—Menten equation? Molecular and Cellular Biochemistry. 1977;15(3):173–8.

